# US Dog Importations during the COVID-19 Pandemic: Do we have an erupting problem?

**DOI:** 10.1101/2021.07.23.453524

**Authors:** Emily G Pieracci, Cara Williams, Ryan M Wallace, Cheryl Kalapura, Clive M Brown

**Affiliations:** Division of Global Migration and Quarantine, National Center for Emerging and Zoonotic Infectious Diseases, CDC; Division of High-Consequence Pathogens and Pathology, National Center for Emerging and Zoonotic Infectious Diseases, CDC; Division of Performance Improvement and Field Services, Center for State, Tribal, Local and Territorial Support, CDC

## Abstract

Dog importation data from 2018-2020 were evaluated to ascertain whether the dog importation patterns in the United States changed during the COVID-19 pandemic, specifically with regard to denial of entry. Dog denial of entry reports from January 1, 2018, to December 31, 2020, stored within the Centers for Disease Control and Prevention (CDC) Quarantine Activity Reporting System, were reviewed. Basic descriptive statistics were used to analyze the data. Reason for denial, country of origin, and month of importation were all examined to determine which countries of origin resulted in the largest number of denials, and whether there was a seasonal change in importations during the COVID-19 pandemic (2020), compared to previous years (2018 and 2019). During 2020, CDC denied entry to 458 dogs. This represents a 52% increase in dogs denied entry compared to the averages in 2018 and 2019. Dogs were primarily denied entry for falsified rabies vaccination certificates (56%). Three countries exported 74% of all dogs denied entry into the United States, suggesting that targeted interventions may be needed for certain countries. Increased attempts to import inadequately vaccinated dogs from countries with canine rabies in 2020 may have been due to the increased demand for domestic pets during the COVID-19 pandemic. Educational messaging should highlight the risk of rabies and the importance of making informed pet purchases from foreign entities to protect pet owners, their families, and the public.

## Introduction

The canine rabies virus variant (CRVV) was eliminated from the United States in 2007, and the Centers for Disease Control and Prevention (CDC) regulates the importation of dogs to prevent the reintroduction of CRVV [1]. Rabies is nearly always fatal once clinical signs begin, and all mammals, including humans, are susceptible to infection [2]. Worldwide, CRVV accounts for 98% of human rabies deaths [2]. The CDC regulates the importation of dogs into the U.S. under 42 CFR 71.51, and in 2019 issued an Order under this authority, due to the public health risk that rabies presents, requiring that dogs imported into the United States from countries with enzootic CRVV (referred to as high-risk countries) to be fully vaccinated against rabies prior to arrival [3,4]. CDC rabies vaccination certificate (RVC) requirements are available to the public on CDC’s website [5]. Dogs from high-risk countries are required to be vaccinated against rabies on or after 12 weeks of age. Additionally, importers must wait 28 days after a dog’s initial vaccination to import the dog into the United States. Therefore, the minimum age at which a dog from a high-risk country can be imported into the United States is 4 months [3,4]. Dogs arriving without valid RVCs are denied entry and returned to the country of origin.

Because all dogs imported into the United States must appear healthy upon arrival, CDC can deny entry and require that dogs receive medical care in the country of origin prior to importation. This requirement is typically only enforced for dogs entering through a land border port of entry; ill dogs that arrive by air are usually referred for local veterinary care because of risks to the dogs and humans during re-exportation. After receiving care from a local U.S. veterinarian and being deemed fit for travel, the dogs are returned to the country of departure.

CDC veterinarians verify the age of dogs arriving at U.S. ports of entry from CRVV high-risk countries to ensure they are at least 4 months of age. A dog’s age can be estimated based on an examination of the dog’s teeth. CDC veterinarians conduct these examinations via photograph, video, or in-person at U.S. ports of entry. CDC follows standard veterinary practice for determining the ages of young dogs [6–8]. The differences between deciduous and permanent teeth are easily discernable in photographs, and the ages at which deciduous and adult dog teeth erupt are well-established in the veterinary profession [6–8].

In early 2020, due to the COVID-19 pandemic, the transportation of animals was significantly reduced by airline carriers worldwide [9]. As Americans transitioned to telework during the pandemic and found themselves at home for a greater proportion of time, people adopted cats and dogs for companionship, and animal shelters were emptied of pets [10–13]. Shelters experienced increased demand for dogs, particularly younger puppies, with some having roughly 200 applications for a single dog [10,13]. When shelters ran out of pets to adopt, potential dog owners looked to rescue organizations, pet stores, and online sellers to find dogs [10]. Low supply and high demand for pets encouraged online puppy sales in the United States and overseas [12]. Online puppy sales saw a spike in demand because many individuals were willing to pay high prices for the immediate companionship of a dog during prolonged periods of isolation and social distancing during the COVID-19 pandemic, and puppies typically garner a higher price compared to older dogs [13,14].

As US residents turn to international puppy supply chains to meet the demand, risks associated with these transactions increasingly result in undesirable complications. Reports of fraudulent sales and scams involving imported dogs are prevalent [11]. People seeking to purchase or adopt animals online are asked to pay transportation and other fees, but often find out that the dog does not meet US entry requirements, does not match the age and description of the dog ordered online, arrives in poor health, or sometimes never existed at all [11,15,16].

Furthermore, increased international dog sales and adoptions also present a public health concern. The US Department of Agriculture regulates breeding of dogs within the United States [17]; however, dogs can be bred and housed under less regulated or non-regulated conditions in many foreign countries. Dogs that are purchased online and arrive from foreign countries may be very young (less than 8 weeks of age), be in poor health or malnourished, lack medical and vaccination history, and carry parasites or communicable diseases of public health concern [18–20]. Additionally, as the authors have witnessed, many dogs arriving from high-risk countries often arrive with falsified, inaccurate or incomplete RVCs, raising questions about the vaccination status of the dog. Well-meaning rescue groups also import large shipments of dogs that unbeknownst to them may be in poor health with subclinical infections that can pose significant public health risks [18]. Three such instances occurred in 2015, 2017, and 2019 when dogs incubating rabies were imported into the United States by rescue groups. In all three cases, the dogs developed clinical rabies shortly after arrival, despite arriving with purported veterinary records indicating the dogs were vaccinated against the disease [21–23].

This evaluation compared dog importation data from 2020 to those of previous years (2018 and 2019) to ascertain whether the dog importation patterns in the United States changed during the COVID-19 pandemic, specifically with regard to denial of entry.

## Methods and Materials

CDC Quarantine Station staff review RVCs for dogs arriving from CRVV high-risk countries. If discrepancies or errors are noted in the RVC, CDC veterinarians are consulted. CDC veterinarians will review the RVC and may examine the dog to verify the age of the dog. CDC veterinarians conduct dental examinations via photograph, video, or in-person at U.S. ports of entry. If a dog does not meet CDC entry requirements a standardized dog denial of entry report is entered by CDC Quarantine Station staff and stored in a secure online database^a^ that records CDC border public health activities including actions taken for CDC-regulated importations at ports of entry. Importers are issued a standardized dog denial of entry letter indicating the reason for the denial of entry. The denial of entry letter, along with all vaccination and medical records presented for the dog, are attached to the report and uploaded into the database. Deidentified data for dog entry denials from January 1, 2018-December 31, 2020, were extracted from the database for inclusion in this analysis. This activity was reviewed by CDC and was conducted consistent with applicable federal law and CDC policy [24]. Monthly data on the intake of dogs into shelters was obtained from the Shelter Animals Count National Database [25].

Traveler count data was requested from U.S. Customs and Border Protection. Summary traveler count information was provided to CDC by month for January 1, 2018-December 31, 2020. The protocol was reviewed and approved by the CDC’s National Center for Emerging and Zoonotic Infectious Diseases Institutional Review Board and was conducted consistent with applicable federal law and CDC policy under 45 C.F.R. part 46, 21 C.F.R. part 56; 42 U.S.C. §241(d); 5 U.S.C. §552a; 44 U.S.C. §3501 et seq.

Basic descriptive statistics were used to analyze the data.^b^ Reason for denial, country of origin, and month of importation were all examined to determine which countries of origin had the highest numbers of denials, and whether there was a seasonal change in importations during the COVID-19 pandemic (2020) compared to previous years (2018, and 2019). Month of importation was included to assess the impact of reduced flights in 2020 and, specifically, the significant reduction in live animal shipments by the airline industry in early 2020 [9].

Dogs arriving from countries that CDC considers high risk for CRVV were included because these dogs are required to have a valid RVC upon arrival [4,26]. Dogs arriving from countries that CDC considers to be low-risk for CRVV or rabies-free were included only if the dogs were ill upon arrival.

Statistical analyses of trends were conducted [27]^b,c^ using a Monte Carlo permutation method to test for significant changes in trends and calculation of the percent change during the trend time period. The number of dog denials and shelter intake of dogs over the 36-month study period were analyzed. Shelter intake was a measure of dog-availability in shelters. Software options were selected for Poisson distribution of standard error. Data were not log-transformed and parameters were set to require trend time periods to contain at least five time points (months). Models were created for six scenarios, reflecting 0-5 allowed trend changes. The model with the lowest p-value was considered most significant and was used for analysis. The resulting model trends reflected the slope of the linear trend line and p-value to indicate if this trend was significantly different from 0%. Additionally, monthly denials and shelter intake data were analyzed.^b^ A logarithmic function was applied to describe the relationship between these two variables, and the resulting model and R^2 value were assessed using quarterly values for both variables.

## Results

During 2020, CDC denied entry to 458 dogs from 29 countries (Table 1). This was a 52% increase in the number of dogs denied entry compared to 2018 (n=304 from 33 countries) and 2019 (n=298 from 20 countries). Three high-risk countries exported 74% (n=338) of all dogs denied entry in the United States: the Russian Federation, Ukraine, and Colombia (Table 2).

**Table 1.**
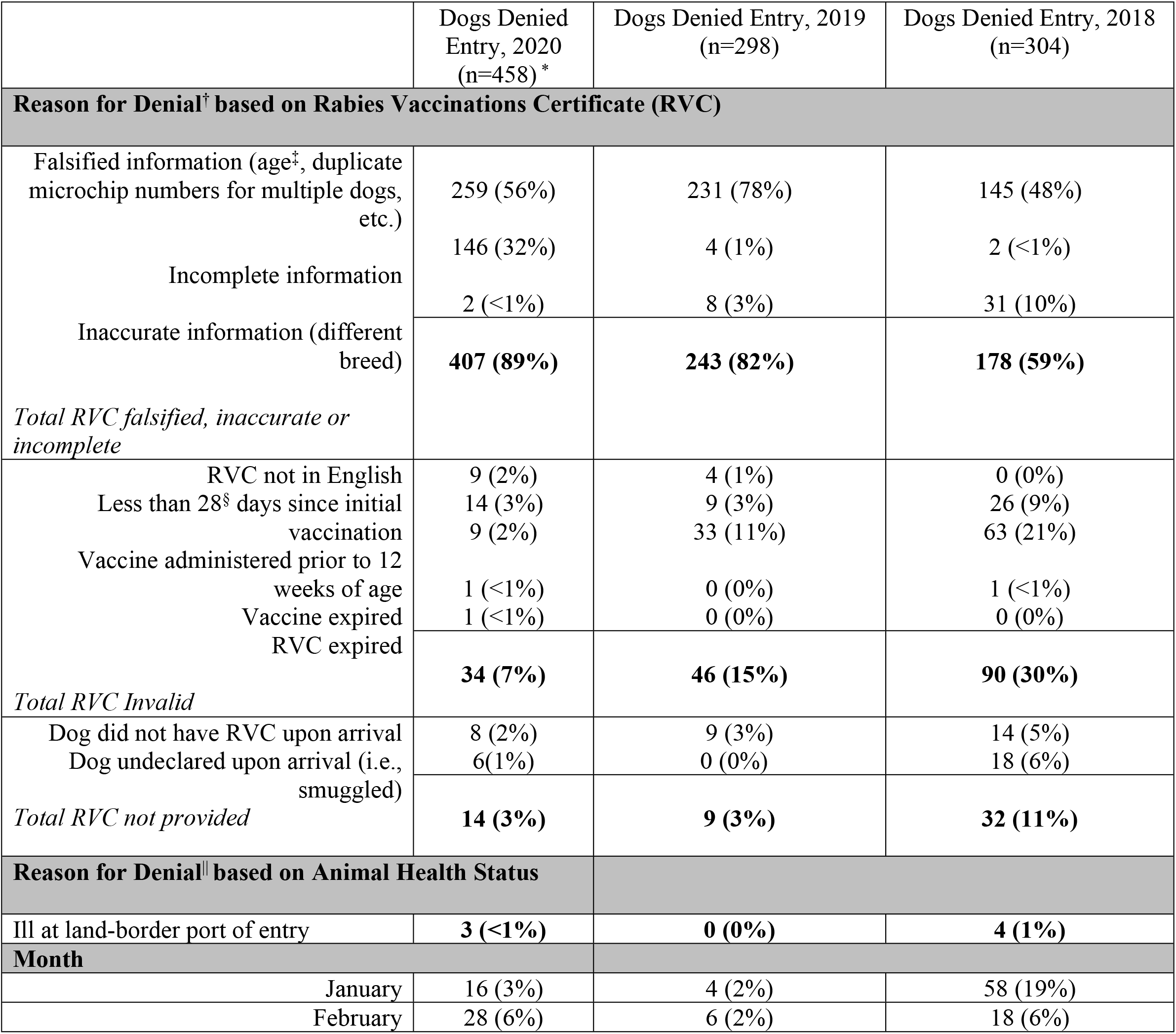
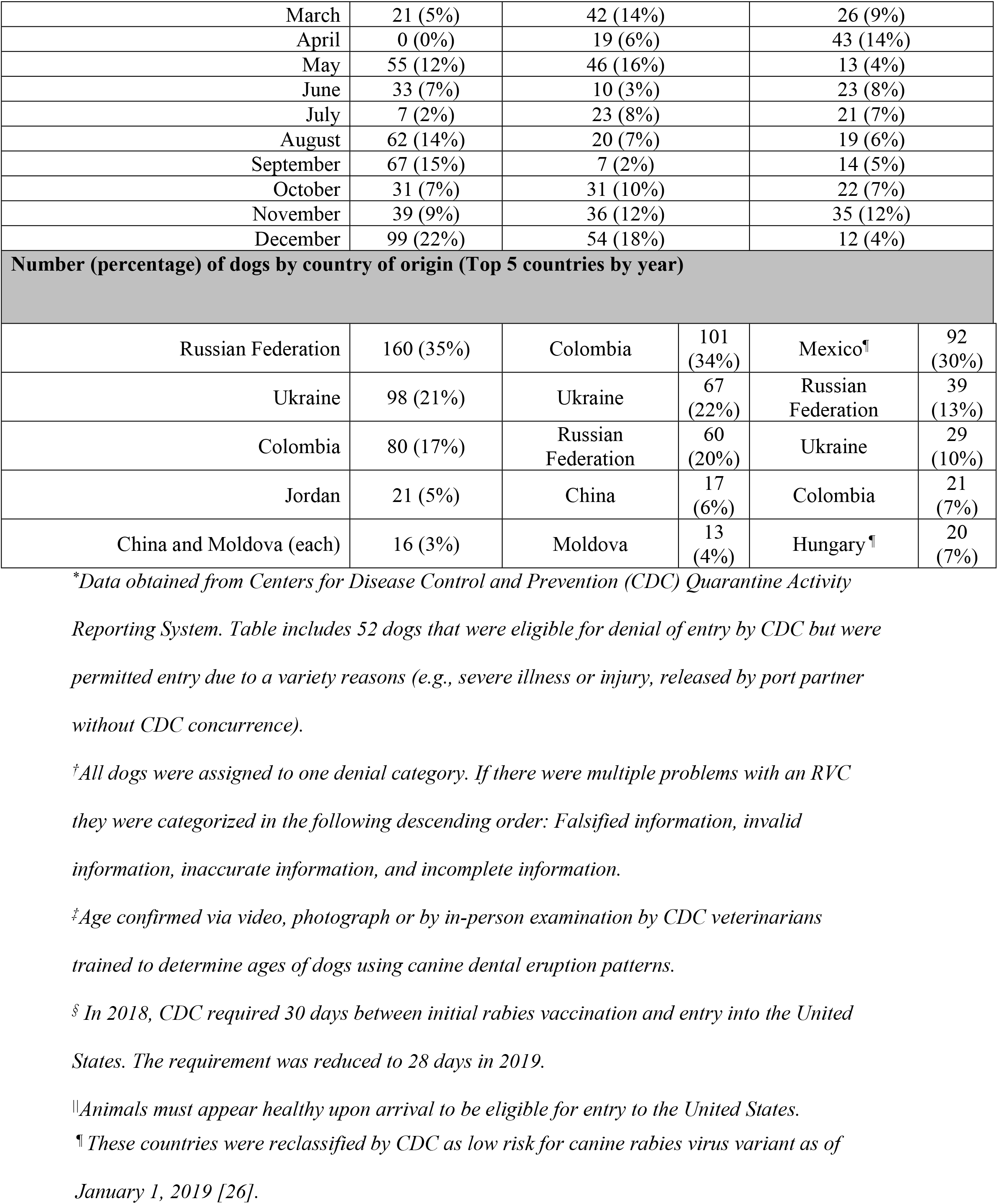
Summary data for dogs denied entry to the United States by year, January 1, 2018 - December 31, 2020.

|  | Dogs Denied Entry, 2020 (n=458) * | Dogs Denied Entry, 2019 (n=298) | Dogs Denied Entry, 2018 (n=304) |
| --- | --- | --- | --- |
| **Reason for Denial<sup>†</sup> based on Rabies Vaccinations Certificate (RVC)** |  |  |  |
| Falsified information (age<sup>‡</sup>, duplicate microchip numbers for multiple dogs, etc.) | 259 (56%) | 231 (78%) | 145 (48%) |
| Incomplete information | 146 (32%) | 4 (1%) | 2 (<1%) |
| Inaccurate information (different breed) | 2 (<1%) | 8 (3%) | 31 (10%) |
| **Total RVC falsified, inaccurate or incomplete** | **407 (89%)** | **243 (82%)** | **178 (59%)** |
| RVC not in English | 9 (2%) | 4 (1%) | 0 (0%) |
| Less than 28<sup>§</sup> days since initial vaccination | 14 (3%) | 9 (3%) | 26 (9%) |
| Vaccine administered prior to 12 weeks of age | 9 (2%) | 33 (11%) | 63 (21%) |
| Vaccine expired | 1 (<1%) | 0 (0%) | 1 (<1%) |
| RVC expired | 1 (<1%) | 0 (0%) | 0 (0%) |
| **Total RVC Invalid** | **34 (7%)** | **46 (15%)** | **90 (30%)** |
| Dog did not have RVC upon arrival | 8 (2%) | 9 (3%) | 14 (5%) |
| Dog undeclared upon arrival (i.e., smuggled) | 6 (1%) | 0 (0%) | 18 (6%) |
| **Total RVC not provided** | **14 (3%)** | **9 (3%)** | **32 (11%)** |
| **Reason for Denial based on Animal Health Status** |  |  |  |
| Ill at land-border port of entry | 3 (<1%) | 0 (0%) | 4 (1%) |
| **Month** |  |  |  |
| January | 16 (3%) | 4 (2%) | 58 (19%) |
| February | 28 (6%) | 6 (2%) | 18 (6%) |
| March | 21 (5%) | 42 (14%) | 26 (9%) |
| April | 0 (0%) | 19 (6%) | 43 (14%) |
| May | 55 (12%) | 46 (16%) | 13 (4%) |
| June | 33 (7%) | 10 (3%) | 23 (8%) |
| July | 7 (2%) | 23 (8%) | 21 (7%) |
| August | 62 (14%) | 20 (7%) | 19 (6%) |
| September | 67 (15%) | 7 (2%) | 14 (5%) |
| October | 31 (7%) | 31 (10%) | 22 (7%) |
| November | 39 (9%) | 36 (12%) | 35 (12%) |
| December | 99 (22%) | 54 (18%) | 12 (4%) |

**Number (percentage) of dogs by country of origin (Top 5 countries by year)**
|  |  |  |  |  |  |
| --- | --- | --- | --- | --- | --- |
| Russian Federation | 160 (35%) | Colombia | 101 (34%) | Mexico<sup>¶</sup> | 92 (30%) |
| Ukraine | 98 (21%) | Ukraine | 67 (22%) | Russian Federation | 39 (13%) |
| Colombia | 80 (17%) | Russian Federation | 60 (20%) | Ukraine | 29 (10%) |
| Jordan | 21 (5%) | China | 17 (6%) | Colombia | 21 (7%) |
| China and Moldova (each) | 16 (3%) | Moldova | 13 (4%) | Hungary<sup>¶</sup> | 20 (7%) |

<sup>\*</sup>Data obtained from Centers for Disease Control and Prevention (CDC) Quarantine Activity
Reporting System. Table includes 52 dogs that were eligible for denial of entry by CDC but were permitted entry due to a variety reasons (e.g., severe illness or injury, released by port partner without CDC concurrence).
<sup>†</sup>All dogs were assigned to one denial category. If there were multiple problems with an RVC they were categorized in the following descending order: Falsified information, invalid information, inaccurate information, and incomplete information.
<sup>‡</sup>Age confirmed via video, photograph or by in-person examination by CDC veterinarians trained to determine ages of dogs using canine dental eruption patterns.
<sup>§</sup> In 2018, CDC required 30 days between initial rabies vaccination and entry into the United States. The requirement was reduced to 28 days in 2019.
<sup>¶</sup>Animals must appear healthy upon arrival to be eligible for entry to the United States.
¶ These countries were reclassified by CDC as low risk for canine rabies virus variant as of January 1, 2019 [26].

**Table 2.**
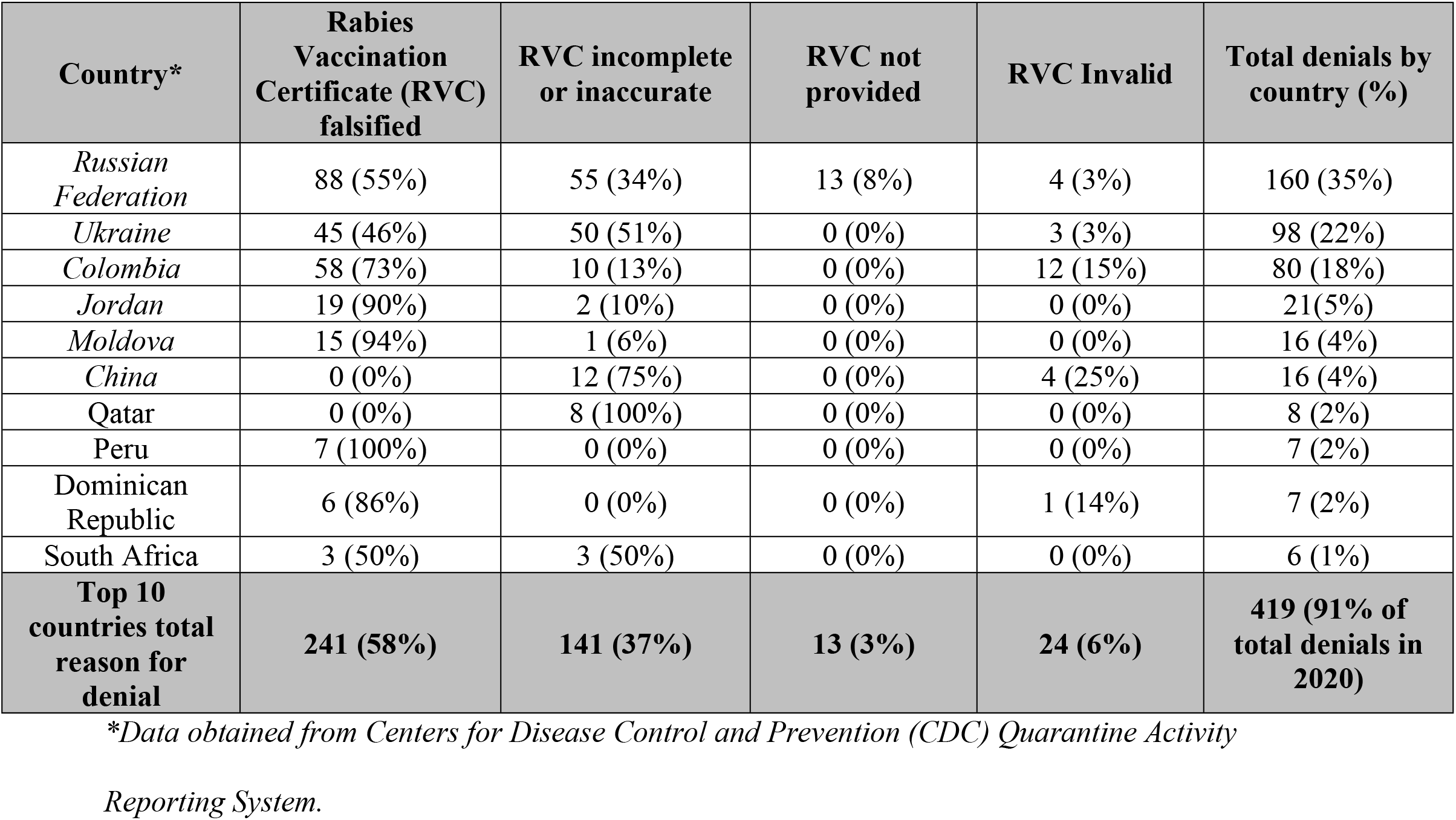
Reasons for dog entry denials by country for the top ten countries of origin, United States, 2020.

Dogs were primarily denied entry for falsified RVCs that documented dogs’ ages as over 4 months when they were determined to be less than 4 months of age by veterinary examination (n=253). Dogs were also frequently denied entry for incomplete information on the RVC (n=146) (Table 1). Fifty-two dogs that were permitted entry despite not meeting CDC entry requirements are included because they were eligible for entry denials. Reasons for their admittance varied: some were accidentally released by federal partners, while some were permitted entry because they were too ill or injured to be returned to the country of origin. For those that were not fully vaccinated on arrival, CDC issued confinement agreements to the importers in coordination with state health officials to ensure the dogs were appropriately vaccinated and confined for a minimum of 28 days after vaccination.

The number and proportion of dogs denied entry for falsified, inaccurate, or incomplete RVCs in 2020 (407, 89%) increased from 2019 (243, 82%) and 2018 (178, 59%).The highest volumes of dog entry denials in 2020 were in December (n=99) and September (n=67). This represented increases of 200% and 538% increase for December and September 2020, respectively, compared to the monthly average for each of those months in 2018 and 2019.

Trend analysis found two significant trend time periods for dog importation denials. Time period one occurred from January 2018 through April 2020, during which the modeled estimate of dog denials declined 89% (p-value 0.051). Time period two occurred from April 2020 through December 2020 and was marked by a significant increase in dog importation denials (224% increase, p-value 0.076). Monthly data for shelter intake of dogs showed three significantly different trend time periods. Time period one occurred from January 2018 – December 2019 and was characterized by stable intake values (5% increase, p-value 0.30). Time period two occurred from December 2019 through April 2020, during which time the intake of dogs in shelters declined 37% (p-value 0.046). The third time period was again characterized by a stable (albeit reduced) intake of dogs, with 8% overall increase (p-value 0.43) (Figure 1). The relationship between dog importation denials and shelter intake of dogs showed a strong, negative, logarithmic association characterized by the following formula (R^2 = 0.49):

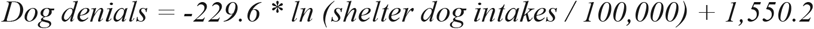

**Figure 1.**
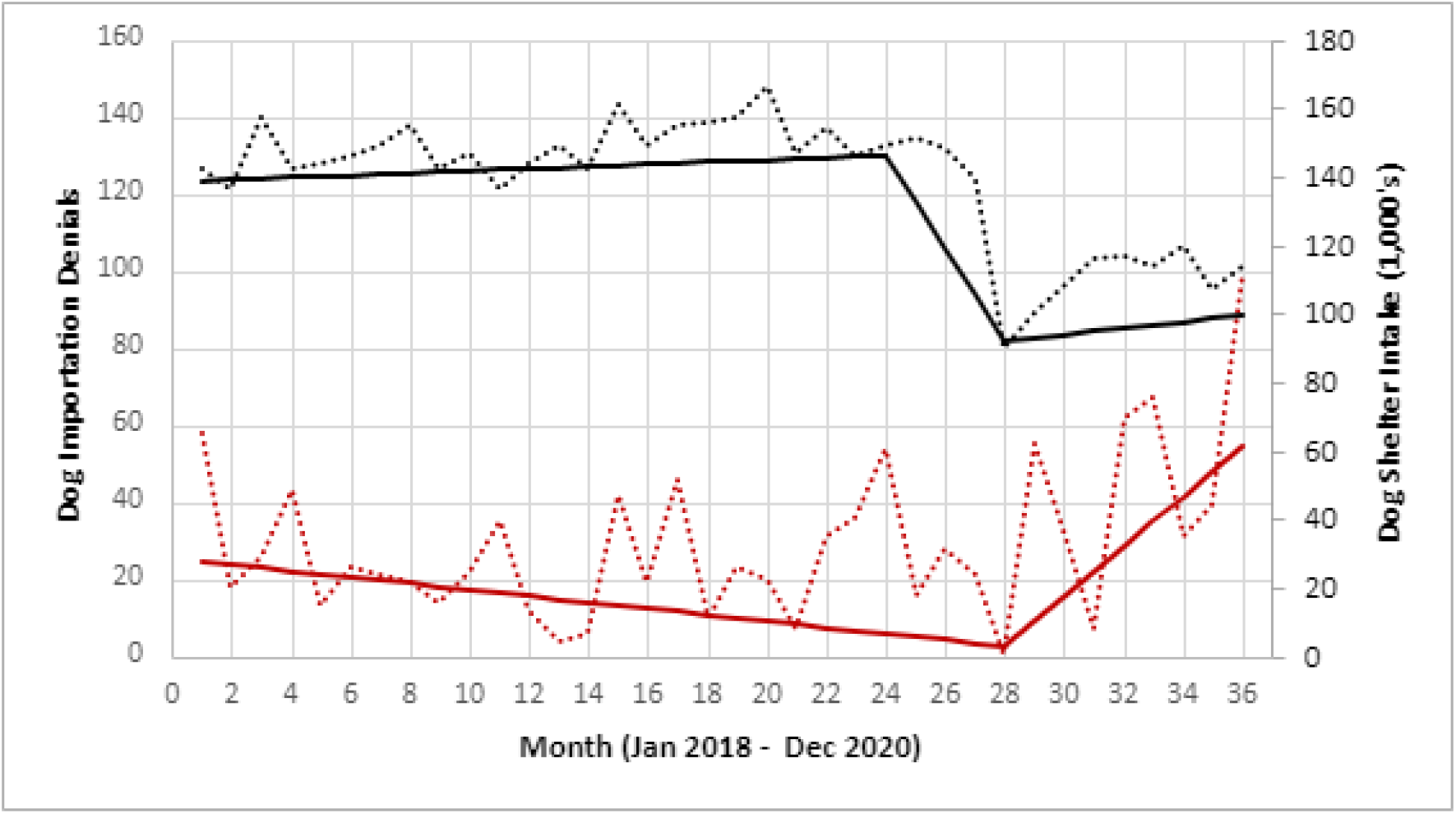
Trends in the intake of dogs in animal shelters compared to dog denials, United States, 2018 – 2020.*^†^. *\* Black dotted line reflects the monthly intake of dogs in shelters. The solid line represents the modeled values from trend analysis and show three trend periods (January 2018 – December 2019), (December 2019 – April 2020), (April 2020 – December 2020)*. ^†^ *Red dotted line reflects the monthly denial of dogs at CDC quarantine stations. The solid red line represents the modeled values from trend analysis and show two trend periods (January 2018 – April 2020) and (April 2020 – December 2020)*.

Trend analysis found five significant trend time periods for travelers arriving in the United States (2018-2020), however, only one significant time period was noted in 2020. Between December 2019 and April 2020, the number of travelers arriving in the United States significantly decreased by 99% (likely due to the COVID-19 pandemic) (p < 0.001). From April 2020-December 2020 the number of arriving travelers began to increase steadily (Figure 2) (p < 0.001). The log regression was not included for further analysis as a decline in passenger travel was unlikely to be directly related to people purchasing dogs from overseas.

**Figure 2.**
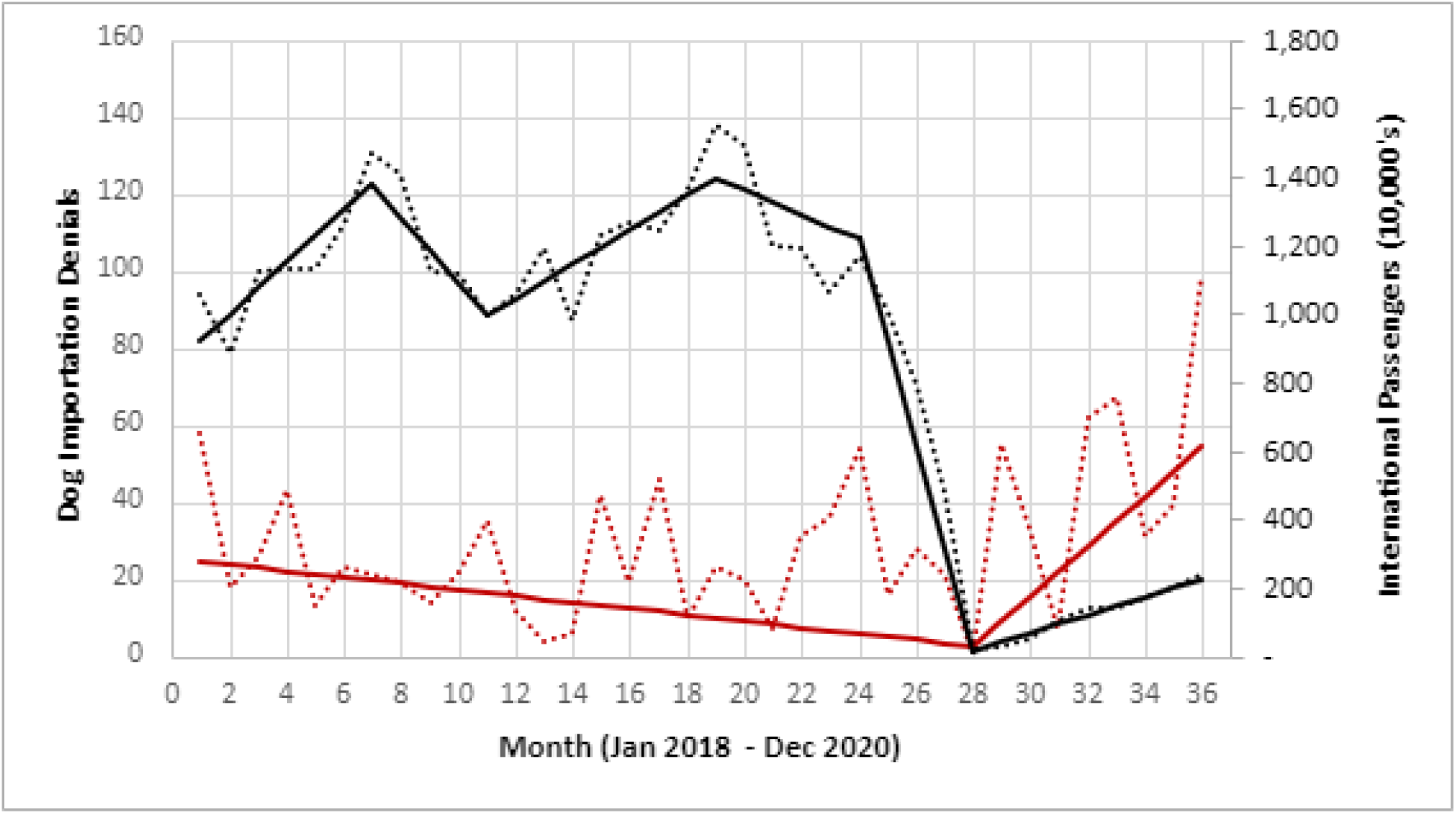
Trends in the number of dogs denied entry and number of travelers arriving into the United States, 2018 – 2020*^†^. *\* Black dotted line reflects the monthly number of travelers arriving in the United States. The solid line represents the modeled values from trend analysis*. ^†^ *Red dotted line reflects the monthly denial of dogs at CDC quarantine stations. The solid red line represents the modeled values from trend analysis and show two trend periods (January 2018 – April 2020) and (April 2020 – December 2020)*.

## Discussion

In early 2020, due to the COVID-19 pandemic, there was a decrease in international flights combined with a temporary suspension of live animal transportation by air carriers. These factors reduced the transportation of animals by airline carriers worldwide [28]. Despite the reduction in flights and animal transports, CDC documented a 52% increase in dog entry denials during 2020 compared to the average number of denials in 2018 and 2019. While the overall number of dogs denied entry for invalid RVCs or missing RVCs appeared to be trending downward from 2018-2020, the number of dogs denied entry for falsified, inaccurate, or incomplete RVCs appears to have increased during this same time period. The increase in falsified, inaccurate, and incomplete rabies vaccination certificates in 2020 may be a result of an increased demand for puppies during the COVID-19 pandemic that led sellers to overlook or attempt to bypass CDC entry requirements. However, similar trends in falsified, inaccurate, or incomplete rabies vaccination certificates were also seen in 2019, suggesting a developing problem with the attempted importation of inadequately vaccinated dogs into the United States. It is important to monitor future trends to determine whether additional interventions are needed to resolve these issues.

The highest volume months of 2020 for dog entry denials were December and September. The December peak may have been due to increased demand of puppies around the holiday season, while the peak in September may have coincided with a return to normal animal transportation policies after the lag seen in early 2020 as a result of COVID-19. A significant change in trend in dog denials occurred in April 2020, characterized by an increase in the monthly rate of dog denials. This trend coincided with in the decline in intake of dogs in shelters. This finding supports a potential association between these two events, marked by a significant reduction in shelter dog populations preceding an increase in fraudulent dog importations by approximately 4 months. The negative logarithmic association between shelter dog intake and dog importation denial had high explanatory value for a univariate comparison. This provides further support that these two events are related, although other factors not assessed in this study also likely have significant influence on dog denial rates.

The attempts to import dogs inadequately vaccinated against rabies during the COVID-19 pandemic is cause for concern as the importation of even one case of canine rabies could result in the re-establishment of CRVV in the United States, which could lead to otherwise preventable deaths among humans, domestic pets, and wildlife. The re-introduction of CRVV in the United States could also lead to sustained transmission in a susceptible host species that could have significant monetary and public health impacts for years to come. There are also health risks to animals and people from possible exposure to an imported rabid dog, including economic costs associated with post exposure prophylaxis and public health investigations.

Additionally, the increase in animal importation challenges faced by federal agencies responsible for protecting public health diverts resources away from other urgent public health efforts, including COVID-19 pandemic response activities. Revision of current CDC dog importation policies should be considered to address this problem. CDC already requires the costs associated with public health mitigation efforts be covered by importers (or the airlines) when dogs do not meet CDC entry requirements. However, throughout 2020, the burden of quarantine, monitoring, veterinary examination, treatment, and revaccination of inadequately vaccinated dogs fell to federal and state public health authorities because non-compliant importers and airline carriers eluded their financial and legal responsibilities. Given the need for a continued COVID-19 response, federal, state, and local public health agencies may not have the resources necessary to address dog importation challenges.

The increase in attempted importations of dogs that did not meet CDC entry requirements during 2020 may have been due to increased demand for pets as people stayed home during the pandemic; lack of availability of animals in shelters due to increased adoptions; enhanced screening by CBP and CDC staff at ports of entry; more permissive or relaxed airline policies in accepting animals for transport after initial restrictions to offset losses from decreased human travel during the pandemic; or importers’ assumption that the public health system infrastructure had deprioritized enforcement of dog entry requirements in favor of COVID-19 related efforts during the pandemic. However, similar patterns documented in 2019 may indicate a developing problem with the attempted importation of inadequately vaccinated dogs in the United States that cannot be explained by the COVID-19 pandemic.

With three countries exporting 74% of dogs denied entry by CDC in 2019 and 2020, targeted interventions are needed to reduce the number of dogs arriving from Russia, Ukraine, and Colombia that do not meet CDC entry requirements. Increased engagement by carriers and brokers to ensure dogs that are transported meet importation country requirements prior to departure could be further supported by industry partners, such as the International Air Transportation Association, which requires air carriers to uphold international transportation requirements when transporting live animals [29].

Finally, educational awareness should be targeted toward animal importers to ensure they understand the risk of rabies and recognize the importance of proper rabies vaccination. Persons wishing to adopt, or purchase pets online should take steps to ensure breeders and rescue organizations are legitimate and ethical. The online purchase of pets has been documented to be challenging and unreliable [9,15] and presents a public health risk as pet owners are unlikely to know their new pet’s medical history, which may include exposure to rabies and other zoonotic diseases of concern. Additionally, attempted imports of dogs under four months of age can put young puppies in danger of serious health problems that may occur during air travel.

There were several limitations in this analysis. First, it is unknown how many dogs enter the United States each year – by land, air, or sea. CDC maintains records of the number of dogs denied entry, their characteristics, and the reason for denial, but the total number of dog importations is unknown. QARS is a passive surveillance system and the number of dogs captured in QARS may not be representative of the true number of dogs entering the United States with falsified, inaccurate, or incomplete rabies vaccination records. Second, the COVID-19 pandemic may have impacted the number and frequency of dog importations in unique ways. As such, the importation data from 2020 may not be representative of dog importation patterns over time. Importation attempts, especially for dogs under four months of age, may be reduced in the future as the demand for new pets decreases and domestic pets become more readily available for adoption or purchase; however, falsified RVCs was a noticeable problem in 2019 prior to the COVID-19 pandemic and the data from 2019 and 2020 could indicate an emerging issue with dog importations. Airline carrier compliance may also improve once more people begin to fly again. The pandemic has had devastating consequences on the airline industry [23], and it is possible some airline carriers sought to offset losses by transporting more animals.

Ultimately, measures to stop the transportation of animals that do not meet US entry requirements should be enforced by air carriers and veterinary officials in the departure country, prior to allowing pets to board long flights that may compromise their health and wellbeing, especially in the case of vulnerable dogs including brachycephalic (snub-nosed) breeds and puppies less than four months of age. It will be important to monitor future trends to determine whether additional interventions are needed to resolve these issues. Consideration of pre-arrival requirements, such as quarantine, booster vaccinations, or serologic tests while in the country of origin may be needed in addition to comprehensive veterinary examinations which should include disease screening and treatment prior to importation when indicated, in order to protect public health and avoid future importations of dogs with rabies or other zoonotic diseases of concern.

## Acknowledgments

We thank Dr. Alida Gertz for her review and feedback of this manuscript and Stephanie Morrison, Melanie Spillane, and Ardath Grills for their assistance with data collection. We thank US Customs and Border Protection for sharing traveler data for the purposes of this analysis. We appreciate the dedication and collaboration of the CDC Quarantine Stations and US Customs and Border Protection in ensuring that dogs presented for importation at US ports of entry meet CDC entry requirements.

## Disclaimer

The findings and conclusions in this report are those of the authors and do not necessarily represent the official position of the Centers for Disease Control and Prevention.

## Footnotes

a Centers for Disease Control and Prevention. *Quarantine Activity Reporting System, version 4.9.8.8.2.2A, 01/25/2021*.

b Microsoft Corporation. *Microsoft Excel 365™*

c Statistical Methodology and Applications Branch, Surveillance Research Program, National Cancer Institute. *Joinpoint Regression Program, Version 4.8.0.1 - April 2020;* Available at: <u>Joinpoint — Joinpoint Help System</u> (cancer.gov). Accessed 5 May 2021.

## Abbreviations List

CBP: Customs and Border Protection
CDC: Centers for Disease Control and Prevention
CFR: Code of Federal Regulations
COVID-19: coronavirus disease 2019
CRVV: canine rabies virus variant
QARS: Quarantine Activity Reporting System
RVC: rabies vaccination certificate

## References

1. Velasco-Villa A, Escobar LE, Sanchez A, et al. Successful strategies implemented towards the elimination of canine rabies in the Western Hemisphere. Antiviral Res 2017; 143: 1–12. https://www.sciencedirect.com/science/article/pii/S0166354216306441?via%3Dihub

2. Hampson K, Coudeville L, Lembo T, et al. Estimating the Global Burden of Endemic Canine Rabies. PLOS Negl Trop Dis. 2015;9(4):e0003709.

3. 42 CFR 71.51. Dogs and Cats. 11 Jan 1985. Available at: <u>SUBPART - Importations</u> (govregs.com) Accessed 10 Apr 2021.

4. Centers for Disease Control and Prevention. Guidance regarding agency interpretation of “rabies-free” as it relates to the importation of dogs into the United States. Federal Register 2019; 84 (21): 724–730. Available at: <u>2019-00506.pdf</u> (govinfo.gov) Accessed 1 Apr 2021.

5. Centers for Disease Control and Prevention. What is a Valid Rabies Vaccination Certificate? Available at: https://www.cdc.gov/importation/bringing-an-animal-into-the-united-states/vaccine-certificate.html. Accessed Apr 20, 2021.

6. Shabestari L, Taylor GN, Angus W. Dental eruption pattern of the beagle. J Dent Res 1967: 276–278.

7. Kremenak CR. Dental exfoliation and eruption chronology in Beagles. J Dent Res 1967; 46 (4): 686–693.

8. Kremenak CR. Dental eruption chronology in dogs: Deciduous tooth gingival emergence. J Dent Res 1969; 48 (6): 1177–84.

9. McMichael C. Nightmare scenarios as travelers attempt to fly with pets amid coronavirus. Available at: <u>Nightmare scenarios as travelers attempt to fly with pets amid coronavirus - ABC News</u> (go.com). Accessed Apr 20, 2021.

10. Kavin K. Dog adoptions and sales soar during the pandemic. The Washington Post. Available at: https://www.washingtonpost.com/nation/2020/08/12/adoptions-dogs-coronavirus/. Accessed 10 Apr 2021.

11. Better Business Bureau. Online puppy scams rising sharply in 2020, BBB warns. Available at: https://www.bbb.org/article/news-releases/23354-bbb-study-update-puppy-scams-rising-in-2020. Accessed 10 Apr 2021.

12. BBC News. Illegal puppy trade warning as sales boom during the Covid pandemic. Available at: https://www.bbc.com/news/uk-scotland-54981798. Accessed 25 Mar 2021.

13. CBC News. Shelters struggle to keep up with skyrocketing demand for pet adoptions during COVID-19. Available at: https://www.cbc.ca/news/canada/british-columbia/high-demand-for-pets-1.5637516. Accessed 5 Apr 2021.

14. Houle MK. Perspective from the field: Illegal puppy imports uncovered at JFK airport. Available at: <u>Illegal Puppy Imports Uncovered | CDC</u>. Accessed 1 Apr 2021.

15. Centers for Disease Control and Prevention. Internet Adoption Scams Involving Imported Pets. Available at: https://www.cdc.gov/importation/internet-scams.html Accessed 25 Mar 2021.

16. Customs and Border Protection. CBP and CDC at LAX Stop Attempt to Smuggle Eight Pomeranian Puppies from Russia. 2020 Aug 24. Available at: https://www.cbp.gov/newsroom/local-media-release/cbp-and-cdc-lax-stop-attempt-smuggle-eight-pomeranian-puppies-russia Accessed 1 Apr 2021.

17. US Department of Agriculture, Animal Plant and Health Inspection Service, Animal Care. Dog Breeder Resource Guide. March 2019. Available at: <u>Dog-Breeder-Resource-Guide.pdf</u> (usda.gov). Accessed 1 Apr 2021.

18. Polak K. Dog transport and infectious disease risk: An international perspective. Vet Clin Small Anim 2019; 49: 599–613.

19. Wagner V, Fernandez-Prada C, Olivier M. A flesh-eating parasite carried by dogs is making its way to North America. The Conversation. 2020 Oct 21. Available at: https://theconversation.com/a-flesh-eating-parasite-carried-by-dogs-is-making-its-way-to-north-america-146652 Accessed 1 Apr 2021.

20. Miller D. The Business of Dog Smuggling and How It Works. Available at: https://humanesocietytampa.org/dog-smuggling-business/ Accessed 25 Mar 2021.

21. Raybern C, Zaldivar A, Tubach S, et. al. Rabies in a Dog Imported from Egypt — Kansas, 2019. Morb Mort Wkly 2020; 69(38): 1374–1377. https://www.cdc.gov/mmwr/volumes/69/wr/mm6938a5.htm

22. Sinclair JR, Wallace RM, Gruszynski K, Freeman MB, Campbell C, Semple S, et al. Rabies in a Dog Imported from Egypt with a Falsified Rabies Vaccination Certificate — Virginia, 2015. Morb Mort Wkly 2015; 64 (49): 1359–1362.

23. Hercules YJ, Bryant NM, Wallace RN, Nelson RM, Palumbo GA, Williams JE, et al. Rabies in a Dog Imported from Egypt — Connecticut, 2017. Morb Mort Wkly 2018;67(50):1388–91.

24. 45 C.F.R. part 46, 21 C.F.R. part 56; 42 U.S.C. §241(d); 5 U.S.C. §552a; 44 U.S.C. §3501 et seq.

25. Shelter Animals Count National Database. Available at: https://www.shelteranimalscount.org/data-dashboards. Accessed 5 May 2021.

26. Centers for Disease Control and Prevention. Rabies vaccine certificate required when coming from these high-risk countries. Available at: <u>Rabies Vaccine Certificate Required When Coming From These High-Risk Countries | Bringing an Animal into U.S. | Importation | CDC</u> Accessed 25 Mar 2021.

27. Kim HJ, Fay MP, Feuer EJ, Midthune DN. Permutation tests for joinpoint regression with applications to cancer rates. Statistics in Medicine 2000; 19:335–351: (correction: 2001;20:655).

28. Josephs L. U.S. airlines’ 2020 losses expected to top $35 billion as pandemic threatens another difficult year. CNBC. Available at: <u>Covid: U.S. airline 2020 losses expected to top $35 billion in dismal year</u> (cnbc.com) Accessed 1 Apr 2021.

29. International Animal Transportation Association. Live Animals. Available at: https://www.iata.org/en/programs/cargo/live-animals/ Accessed 1 Apr 2021.

